## Supplementary Information for "Neural attentional filters and behavioural outcome follow independent individual trajectories over the adult life span"

### for

### Supplemental figures

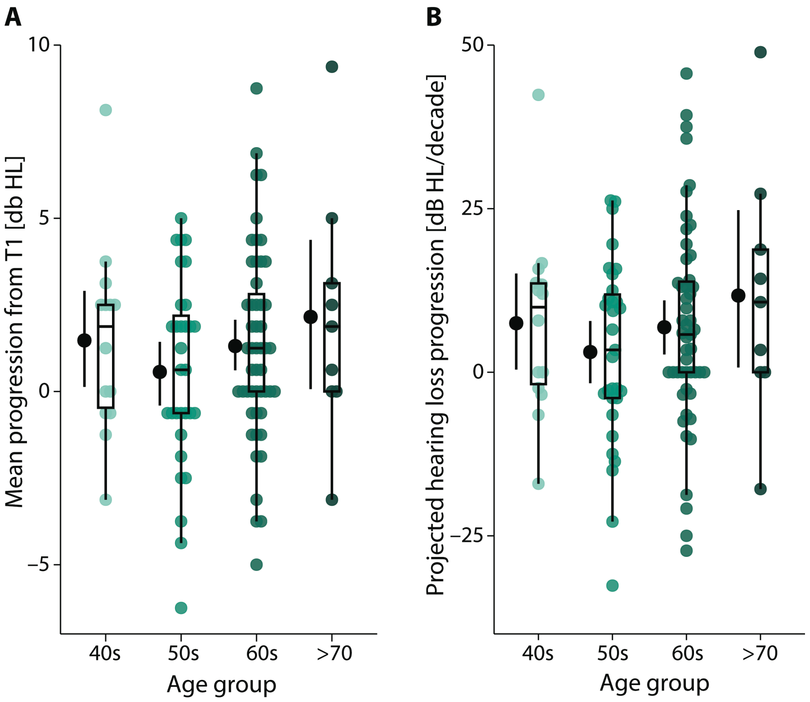


**Fig S1. Observed and projected hearing loss progression. A,** Mean progression of hearing loss from T1 to T2 as calculated from two-year differences in the individual pure-tone average hearing thresholds at 0.5, 1,2, and 4 kHz. Coloured dots represent the single-subject difference across four age groups (total N = 105 participants), black dots and error bars indicate grand-average and bootstrapped 95% CI. Box plots show median centre line, 25th to 75th percentile hinges, whiskers indicate minimum and maximum within 1.5 × interquartile range. **B**, analogue to panel A but showing the projected hearing loss progression [increase in db HL per decade] assuming a constant rate of change per individual.

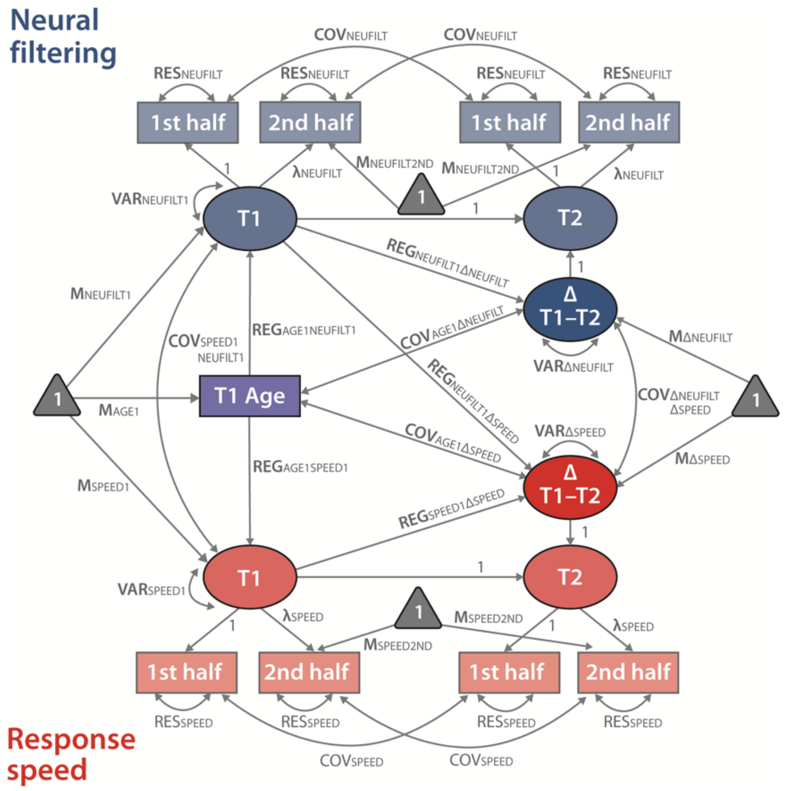


**Fig S2**. **Full bivariate latent change score model of response speed and neural filtering**. Model estimated brain–behaviour baseline–change (via regression) and change–change (via their covariance) relationship. First order latent variables (T1, T2) were constructed from manifest neural and behavioural metrics extracted from the first and second half of the experiment, respectively. M, mean; COV, covariance; REG, regression path; RES, residual; **λ**, factor loading; Δ, change. Residuals and covariances of manifest variables of the same metric were each set to be equal.

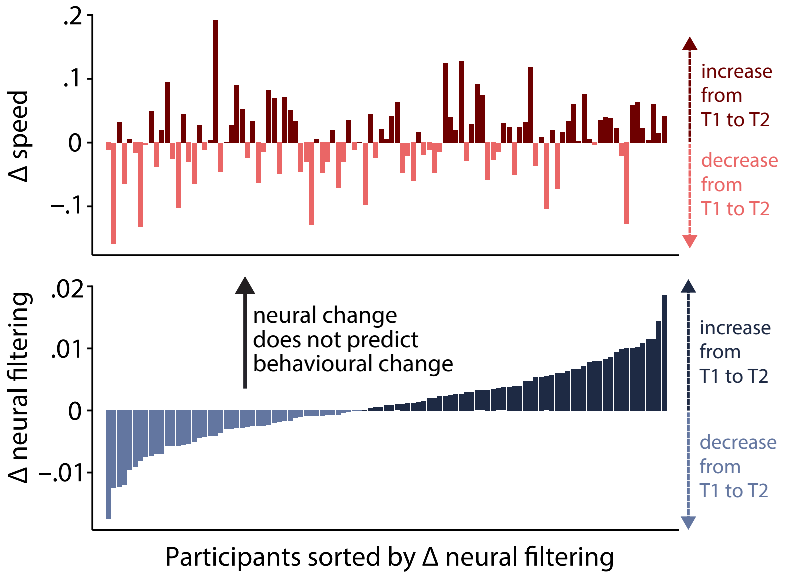


**Fig S3**. **Change in neural filtering does not predict change in response speed.** Bottom, Individuals sorted by their change in neural filtering. Positive values indicating an increase in neural filtering over time, negative values indicating a decrease. Applying the same ordering to individuals' longitudinal change in response speed (top) does not uncover a systematic relationship in cross-domain changes.

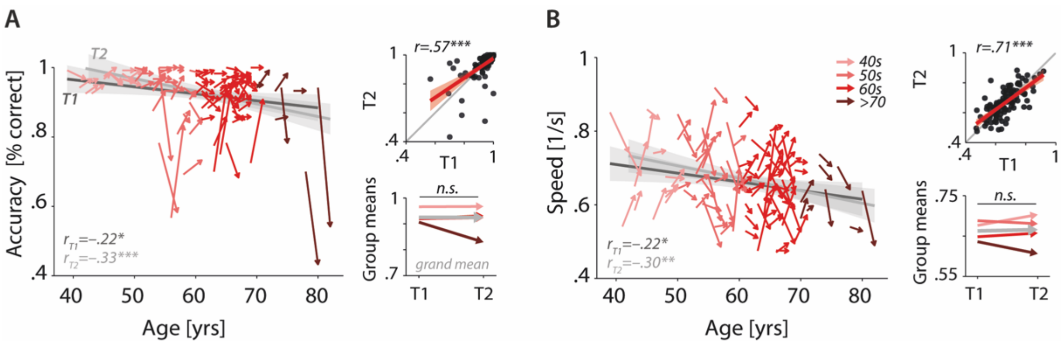


**Fig S4. Cross-sectional and longitudinal change in accuracy (A) and response speed (B) averaged across selective-attention trials, only.** Coloured vectors (colour-coding four age groups for illustrative purposes, only) in the left subpanels show individual T1–T2 change along with the cross-sectional trend plus 95% confidence interval (CI) separately for T1 (dark grey) and T2 (light grey). Top right, correlation of T1 and T2 as measure of test-retest reliability along with the 45° line (grey) and individual data points (black circles). Bottom right, mean longitudinal change per age group and grand mean change (grey).

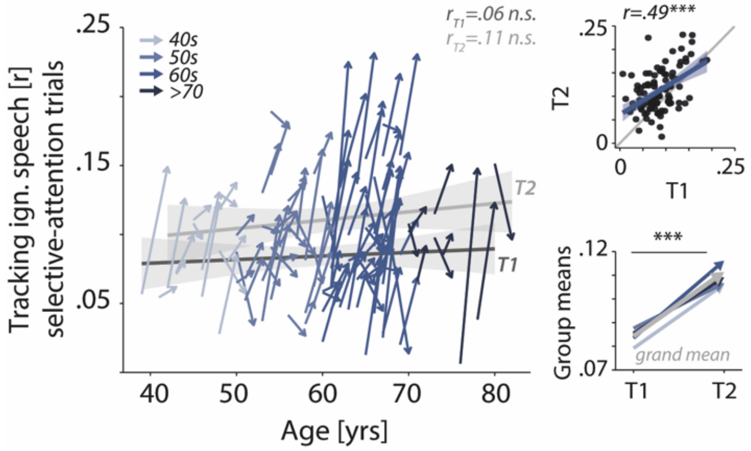


**Fig S5. Cross-sectional and longitudinal change in neural tracking of ignored speech averaged across selective-attention trials, only.** Coloured vectors (colour-coding four age groups for illustrative purposes, only) in the left subpanels show individual T1–T2 change along with the cross-sectional trend plus 95% confidence interval (CI) separately for T1 (dark grey) and T2 (light grey). Top right, correlation of T1 and T2 as measure of test-retest reliability along with the 45° line (grey) and individual data points (black circles). Bottom right, mean longitudinal change per age group and grand mean change (grey).

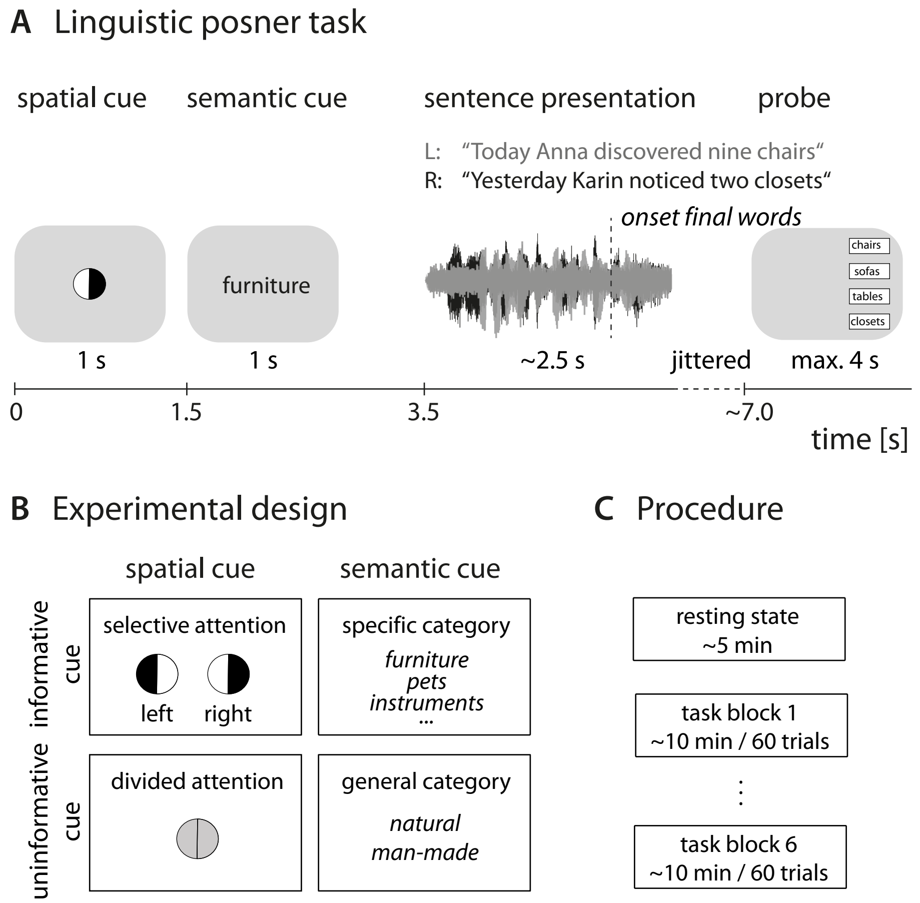


**Figure S6. Experimental design and procedure.** A, Trial-structure of dichotic listening task. In each trial participants listened to two dichotically presented short sentences spoken by the same speaker. They had to identify the final word in one of the two sentences. Sentence presentation was preceded by two visual cues. B, 2x2 design in which the presentation of the spatial-attention and semantic cue was fully crossed. Each cue was presented in either an informative or uninformative neural version. C, Prior to the listening task, we recorded resting-state EEG data. The listening task was presented in 6 blocks of 60 trials with break in between.
